# Age-related differences in negative cognitive empathy but similarities in positive affective empathy

**DOI:** 10.1101/2020.04.03.024877

**Authors:** Maryam Ziaei, Lena Oestreich, David C. Reutens, Natalie C. Ebner

## Abstract

Empathy, among other social-cognitive processes, changes across adulthood. More specifically, cognitive components of empathy (understanding another’s perspective) appear to decline with age, while findings for affective empathy (sharing another’s emotional state) are rather mixed. Structural and functional correlates underlying cognitive and affective empathy in aging and the extent to which valence affects empathic response in brain and behavior are not well understood yet. To fill these research gaps, younger and older adults completed a modified version of the Multifaceted Empathy Test, which measures both cognitive and affective empathy as well as empathic responding to both positive and negative stimuli (i.e., positive vs. negative empathy). Adopting a multimodal imaging approach and applying multivariate analysis, the study found that for cognitive empathy to negative emotions, regions of the salience network including the anterior insula and anterior cingulate were more involved in older than younger adults. For affective empathy to positive emotions, in contrast, younger and older adults recruited a similar brain network including main nodes of the default mode network. Additionally, increased structural integrity (fractional anisotropy values) of the posterior, but not the anterior, cingulum bundle was related to activation of default mode regions during affective empathy for positive emotions in both age groups. These findings provide novel insights into the functional networks subserving cognitive and affective empathy in younger and older adults and highlight the importance of considering valence in empathic response in aging research. Further this study, for the first time, underscores the role of the posterior cingulum bundle in higher-order social-cognitive processes such as empathy, specifically for positive emotions, in aging.

## Introduction

Mounting evidence suggests age-related changes in social-cognitive capacities including perception of eye gaze (Slessor et al., 2010; Ziaei et al., 2016), emotional facial expression recognition (Ruffman et al., 2008), and theory of mind (Henry et al., 2013). Compared to other social-cognitive components, empathy has received considerably less attention in aging research. Furthermore, the few existing studies on age-related differences in empathic response have almost exclusively used self-report and have captured more trait-like aspects of the construct (e.g., using the Interpersonal Reactivity Index; Davis, 1983). These previous studies largely agree that aging is associated with decline in cognitive empathy, the ability to decode and understand another’s perspective (Beadle & de la Vega, 2019; Henry et al., 2013). However, evidence is less clear for affective empathy, i.e., the affective sharing of another’s emotional state (Singer & Lamm, 2009), for which some studies suggest no age-related differences (Bailey et al., 2008; Beadle et al., 2012) while other studies support an increase with age (Grühn et al., 2008; O’Brien et al., 2012).

Rather sparse are studies measuring state empathy, specifically in aging, for example by using experimentally induced alterations in the state of affective empathy. In particular, Sze and colleagues (2012) manipulated empathic response by showing uplifting or distressing films and reported an age-related linear increase in empathic concern and personal distress in response to both types of films. Similarly, Bailey et al. (2018) found age-related increased emotional distress and reactivity towards another’s pain; and this enhanced affective empathy predicted prosociality (i.e., willingness to help). Finally, Beadle et al. (2015) did not find evidence for age-group differences in affective empathy or personal distress in response to cancer patients describing their experiences with the disease. This currently limited and mixed knowledge base on age effects in cognitive and affective empathy warrants additional research (see also Bailey et al., 2021; Ebner et al., 2017, for overviews).

Over the last two decades robust evidence suggests prioritized processing of positive over negative information among older (compared to younger) adults. This effect is reflected in greater attention to, and memory for, positive over negative stimuli and has been termed the “positivity effect” in aging (Mather & Carstensen, 2003; Reed & Carstensen, 2012; Ziaei & Fischer, 2016; Ziaei, Salami, et al., 2017; Ziaei et al., 2015). The positivity effect has been interpreted as a motivational, goal-oriented shift with age. Closely related to this notion of a positivity effect in aging is the motivational empathy account, also referred to as ‘empathic concern’ (Weisz & Zaki, 2018). According to this account, an observers’ motivation/social goals can manifest in the reduction or intensification of empathy towards a target.

To date, understanding of the impact of valence on empathy in aging is still very limited. First findings in younger adults support the distinction of empathic response to positive vs. negative stimuli, known as positive vs. negative empathy, respectively (Morelli et al., 2015). Previous work suggests that people use the emotion expressed by others as a social signal to interpret what others are feeling (Van Kleef, 2009). Thus, based on robust evidence of an age-related shift in processing positive over negative information (i.e., positivity effect), and in line with the motivational account of empathy, it is reasonable to assume that older adults’ empathic response is particularly impacted by valence of to-be-processed information.

The anterior insula, mid and dorsal anterior cingulate gyrus, and temporo-parietal junction have been identified as key brain regions involved in empathy (Bernhardt & Singer, 2012; Bzdok et al., 2012b; Decety & Jackson, 2006; Lamm et al., 2011). However, to date, only two studies have investigated the neural substrates underlying empathy in older adults. In particular, Chen and colleagues (2014) asked younger and older adults to rate their feelings towards another’s pain and found an age-related decrease in activation of insula and anterior cingulate during this task. Additionally, Riva et al. (2018) reported reduced insula activity during both pleasant and unpleasant touch using a visuotactile stimulation paradigm among older female participants.

While meta-analyses support the importance of insula and anterior cingulate regions with a core “empathy network”, a distinction between affective-perceptual vs. cognitive-evaluative empathy networks has been proposed (Bellucci et al., 2020; Fan et al., 2011). In this context, the anterior insula has been suggested to be involved in affective-perceptual empathy and the anterior cingulate in cognitive-evaluative empathy. Furthermore, as empathy is a complex and multidimensional process, it is likely that empathic response activates large-scale brain networks and not just individual regions, but a multivariate approach to the study of empathy has not been undertaken yet.

White matter tracts, such as the cingulum bundle, which links the frontal lobe with the precuneus, posterior cingulate cortex, hippocampus, and parahippocampus (Wakana et al., 2004), are believed to be critical in attention, memory, executive functioning, and emotional processing (Keedwell et al., 2016; van den Heuvel et al., 2008; Wu et al., 2016). Also, more integrity in the anterior subdivision of the cingulum bundle was associated with better cognitive control in older adults (Metzler-Baddeley et al., 2012). However, structural pathways that subserve empathic responding have not been well investigated yet, and currently unknown is the extent to which integrity of the anterior and the posterior cingulum bundle may facilitate higher-order social-cognitive processes, such as empathy, among older adults.

To address the above-identified research gaps and to integrate previously parallel lines of work, the present study had three major aims: to examine the extent to which functional activation involved in cognitive and affective empathy: *(i)* differed between younger and older adults; *(ii)* in interaction with stimulus valence; and *(iii)* was related to cingulum bundle microstructure. Here, younger and older adults completed a modified version of the Multifaceted Empathy Test (MET) (Dziobek et al., 2011) that comprised both cognitive and affective empathy components (as well as a neutral age perception control condition) and included positive, negative, and neutral images to allow for a systematic investigation of valence effects on empathic response (Mazza et al., 2015). Both structural and functional brain images were acquired to determine the association between functional network activation during the empathy task and white matter tract integrity known to play a role in emotional processing and aging.

We hypothesized that older relative to younger adults would show poorer performance, and that the age groups would display differential recruitment of brain networks (e.g., the limbic system; Yu & Chou, 2018), during cognitive empathy (*Hypothesis 1a*; Beadle & de la Vega, 2019). In contrast, we predicted that performance during affective empathy would be comparable between younger and older adults (Beadle & de la Vega, 2019) and that engagement of affective empathy-related brain regions (such as the insula and the anterior cingulate cortex; Singer & Lamm, 2009) would be comparable between the two age groups (*Hypothesis 1b*).

Furthermore, we expected that the age groups would differ in their recruitment of brain networks in response to negative and positive stimuli, with reduced activity of the salience network for negative (*Hypothesis 2a*) but enhanced or equal activity of the default mode network for positive (*Hypothesis 2b*) stimuli among older (relative to younger) participants. These predictions were based on evidence that positive and negative empathy selectively activate regions associated with positive (e.g., ventromedial prefrontal cortex) and negative (e.g., anterior insula, dorsal anterior cingulate cortex) affect, respectively (Morelli et al., 2015; Ziaei et al., 2019). We have previously demonstrated that positive affect was associated with activation in regions within the default mode network in older adults (Ziaei, Ebner, et al., 2017). To our knowledge, however, no studies have directly compared neural correlates of positive vs. negative stimuli between younger and older adults during the empathy task. Our predictions built on research in emotion recognition that suggests that regions such as the ventromedial prefrontal cortex and posterior cingulate cortex are involved in processing positive emotions, while regions such as the insula and anterior cingulate are involved in processing negative emotions. Here we also explored age-related differences in the anterior and posterior subdivision of the cingulum bundle in their association with functional activation during both cognitive and affective empathy. We anticipated that higher fractional anisotropy of the anterior cingulum bundle would be related to functional activation during cognitive empathy and posterior cingulum would be related to functional activation during affective empathy among older adults.

## Method

### Participants

Twenty-six younger (18-25 years of age) and 25 older (65-80 years of age) adults participated in this study. Due to large head movement (> 1.5mm), one younger and one older adult were excluded, leaving 25 younger (*M* = 21.72, *SD* = 3.81; 13 females) and 24 older (*M* = 71.75, *SD* = 3.70; 14 females) participants for final brain imaging data analysis. All younger participants were University of Queensland undergraduate students who were reimbursed either with course credits or AUD$20 per hour. Older participants were volunteers from the community, recruited through advertising in public notice boards of different clubs, libraries, churches, and the University of Queensland’s Aging Mind Initiative. Older participants were reimbursed with AUD$20 per hour.

All participants were right-handed, English speakers who had normal or corrected-to-normal vision using magnetic resonance imaging (MRI) compatible glasses, and no history of diagnosed psychiatric illnesses, no history of medication for psychiatric illnesses, no cardiovascular disease, head/heart surgery, or neurological impairment (e.g., epilepsy). The age groups were comparable in years of education and gender distribution (Table 1). All older adults scored above the recommended cut-off of 27 (*M* = 28.76, *SD* = 1.26) on the Mini Mental State Examination (Folstein et al., 1975).

**Table 1.** Descriptive data (means and standard deviations) and inferential statistics for background measures in younger and older adults

|  | Younger Adults |  | Older Adults |  | Age-Group Differences |  |
| --- | --- | --- | --- | --- | --- | --- |
|  | *M* | *SD* | *M* | *SD* | *t* | *df* |
| **Demographics** |  |  |  |  |  |  |
| Age (years) | 21.72 | 3.81 | 71.75 | 3.70 | 46.52** | 47 |
| Gender | 13 females |  | 14 females |  |  |  |
| Education (years) | 15.50 | 2.46 | 17.47 | 8.95 | 1.06 | 47 |
| **Executive Functioning** |  |  |  |  |  |  |
| Verbal Fluency Test | 47.84 | 10.67 | 53.33 | 12.67 | 1.64 | 47 |
| Stroop Interference Score | 0.23 | 0.25 | 0.28 | 0.19 | 0.73 | 47 |
| Task Switching Index | 20.83 | 8.64 | 24.46 | 12.08 | 1.21 | 47 |
| Raven IQ | 6.40 | 1.77 | 5.66 | 2.01 | 1.35 | 47 |
| **Emotional Well-Being** |  |  |  |  |  |  |
| DASS-21 |  |  |  |  |  |  |
| *Stress* | 6.88 | 6.63 | 1.33 | 2.33 | 3.87** | 47 |
| *Anxiety* | 5.12 | 3.60 | 1.00 | 1.56 | 5.14** | 47 |
| *Depression* | 10.40 | 5.94 | 3.66 | 3.85 | 4.68** | 47 |
| **Empathy** |  |  |  |  |  |  |
| IRI total | 66.04 | 10.21 | 61.00 | 12.46 | 1.55 | 47 |
| *Fantasy* | 16.12 | 4.70 | 13.75 | 5.51 | 1.62 | 47 |
| *Empathic concern* | 17.88 | 3.63 | 18.54 | 3.56 | 0.64 | 47 |
| *Perspective taking* | 19.44 | 4.11 | 19.62 | 3.51 | 0.16 | 47 |
| *Personal distress* | 12.04 | 5.67 | 8.41 | 4.26 | 2.5* | 47 |
| Empathy Quotient | 44.44 | 11.16 | 48.95 | 11.68 | 1.38 | 47 |
| **RMET** | 25.76 | 4.20 | 27.95 | 4.29 | 1.80 | 47 |

*Notes.* \* $p < 0.05$ , \*\* $p < .001$ . Verbal Fluency Test (Benton & Hamsher, 1976): total number of words for letters F, A, and S; Stroop Interference Score (Ziaei et al., 2015) = ((response time in incongruent trials – response time in neutral trials)/response time in neutral trials); Task-Switching Index (Reitan & Wolfson, 1986): Trail Making Test Part B – Trail Making Test Part A; Raven IQ (Kirchner, 1958) = total number of correct responses; DASS-21 = Depression Anxiety Stress Scale (Lovibond & Lovibond, 1995); IRI = Interpersonal Reactivity Index (Davis, 1983); RMET = Reading the Mind in the Eyes Test (Baron-Cohen et al., 2001); $df$ = degree of freedom; $M$ = mean; $SD$ = standard deviation; $t$ = student t-test.

**Table 2.**
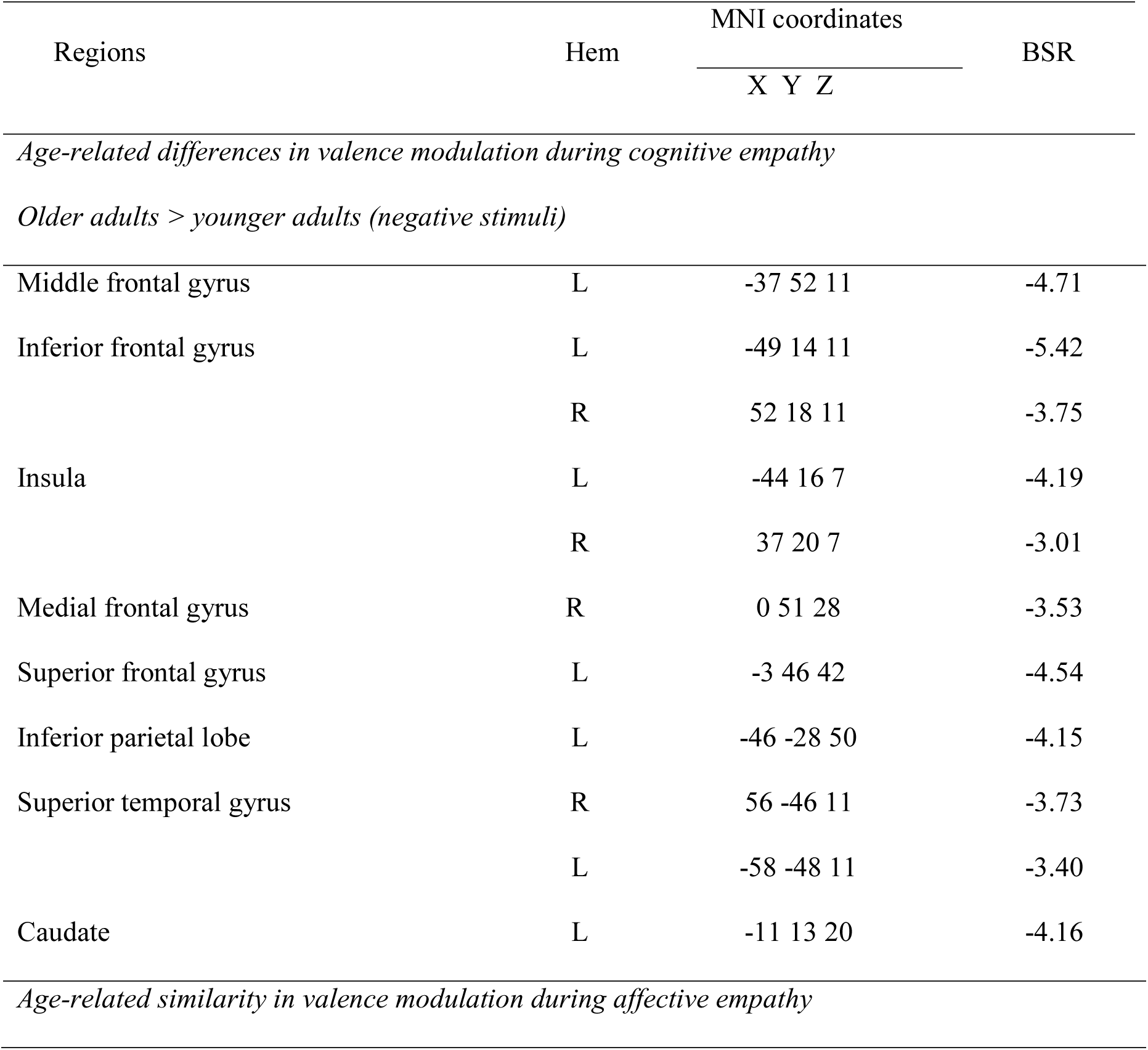
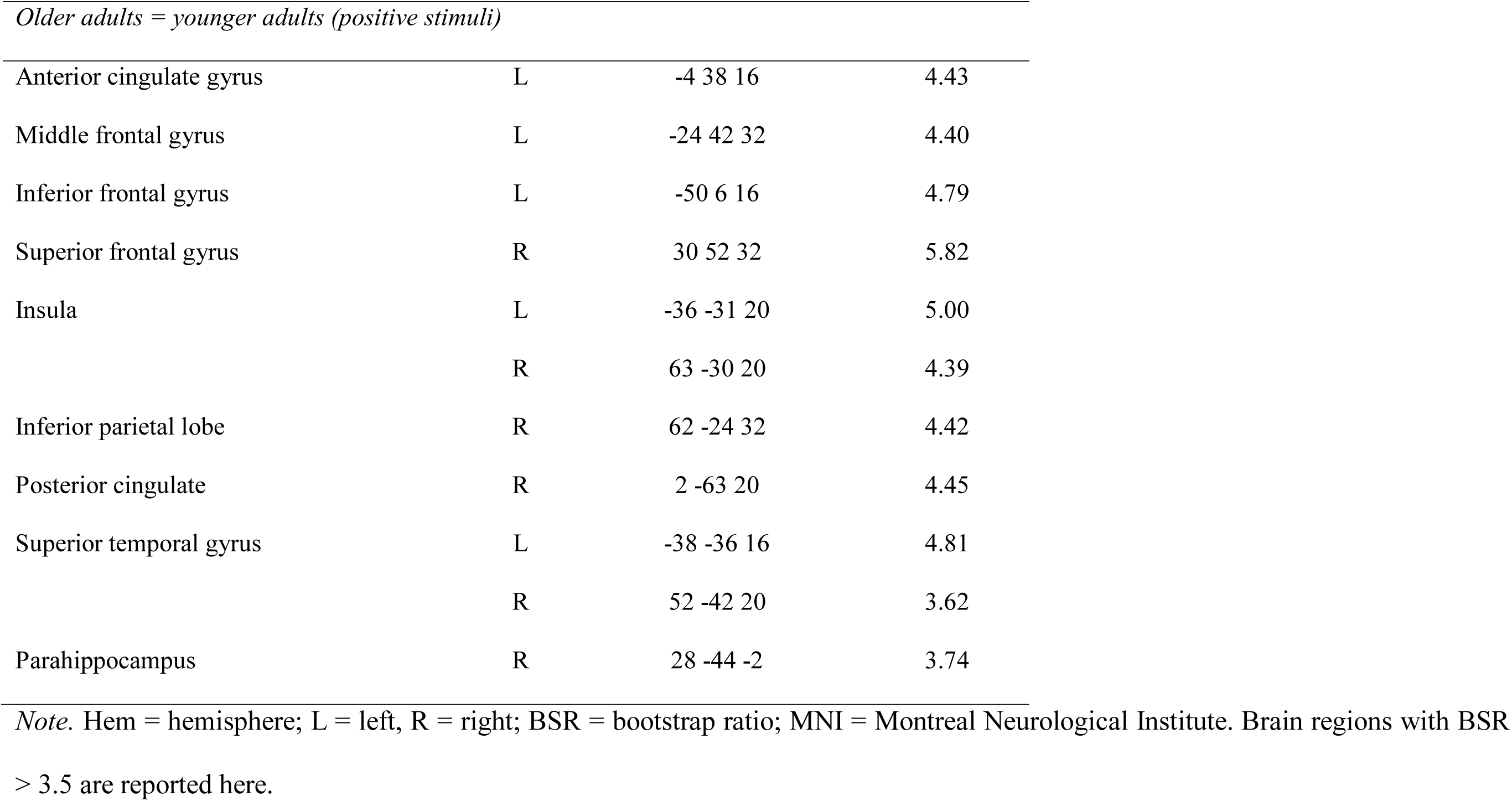
Whole-brain results showing pattern of activity across younger and older adults by valence modulation (i.e., positive and negative stimuli)

| Regions | Hem | MNI coordinates | BSR |
| --- | --- | --- | --- |
|  |  | X Y Z |  |
| *Age-related differences in valence modulation during cognitive empathy* |  |  |  |
| *Older adults > younger adults (negative stimuli)* |  |  |  |
| Middle frontal gyrus | L | -37 52 11 | -4.71 |
| Inferior frontal gyrus | L | -49 14 11 | -5.42 |
|  | R | 52 18 11 | -3.75 |
| Insula | L | -44 16 7 | -4.19 |
|  | R | 37 20 7 | -3.01 |
| Medial frontal gyrus | R | 0 51 28 | -3.53 |
| Superior frontal gyrus | L | -3 46 42 | -4.54 |
| Inferior parietal lobe | L | -46 -28 50 | -4.15 |
| Superior temporal gyrus | R | 56 -46 11 | -3.73 |
|  | L | -58 -48 11 | -3.40 |
| Caudate | L | -11 13 20 | -4.16 |
| *Age-related similarity in valence modulation during affective empathy* |  |  |  |
| *Older adults = younger adults (positive stimuli)* |  |  |  |
| Anterior cingulate gyrus | L | -4 38 16 | 4.43 |
| Middle frontal gyrus | L | -24 42 32 | 4.40 |
| Inferior frontal gyrus | L | -50 6 16 | 4.79 |
| Superior frontal gyrus | R | 30 52 32 | 5.82 |
| Insula | L | -36 -31 20 | 5.00 |
|  | R | 63 -30 20 | 4.39 |
| Inferior parietal lobe | R | 62 -24 32 | 4.42 |
| Posterior cingulate | R | 2 -63 20 | 4.45 |
| Superior temporal gyrus | L | -38 -36 16 | 4.81 |
|  | R | 52 -42 20 | 3.62 |
| Parahippocampus | R | 28 -44 -2 | 3.74 |

*Note.* Hem = hemisphere; L = left, R = right; BSR = bootstrap ratio; MNI = Montreal Neurological Institute. Brain regions with BSR > 3.5 are reported here.

### Procedure

The experiment was approved by the Royal Brisbane and Women’s Hospital and the University of Queensland Research Ethics Committees. All participants provided written informed consent prior to enrollment. The study comprised a 1-hour MRI session, followed by a 2-hour behavioral/neuropsychological assessment on the same day. Prior to the MRI, participants received verbal instruction about the MET (described next) and worked on practice trials for familiarization with the trial timing and task sequence. After the MRI, participants completed a series of background measures (described below), were debriefed, and received reimbursement.

### Multifaceted Empathy Task (MET)

We used a modified version of the MET (Dziobek et al., 2011). As depicted in Figure 1, the MET consists of naturalistic images of human faces. The task comprised three experimental conditions: cognitive empathy, affective empathy, and an age perception control. For the cognitive empathy condition, participants were asked to identify “what kind of emotion does this person feel?” by choosing “positive”, “negative”, or “neutral.” For the affective empathy condition, participants were asked to think about their feeling towards the person depicted and rate “how strong is the emotion you feel about this person?” by choosing “low”, “average”, and “high”. This question aimed at evoking emotional responses to the depicted person rather than inferring the emotion experienced by that person (as in the cognitive empathy condition). For the age perception control condition, participants were asked to identify “what is the age of this person?” depicted on the picture by choosing “child”, “adult”, and “elderly”. This condition was used to control for higher-order cognitive processing involved in evaluating the specific stimuli used in this task. Responses were recorded using three keys on an MRI-compatible response box. To reduce working memory load, response options were presented on the screen.

Following Mazza et al. (2015), positive (happy), negative (sad, angry), or neutral faces were presented in each of the three experimental conditions. In particular, we included seven pictures of the same valence in each of the three experimental conditions, with each image presented for six seconds, for a total duration of 42 seconds per block. Compared to the original task (Mazza et al., 2015), we shortened block length from 70 to 42 seconds to improve brain signal (Huettel et al., 2014). Pictures were selected from the original MET and supplemented by pictures from the International Affective Picture System (Lang et al., 2008). The valence and arousal ratings of pictures used in this study were as follow: Negative pictures (valence: *M* = 3.14, *SD* = 1.47; arousal: *M* = 4.78, *SD* = 2.12; e.g., homeless man, crying baby, angry man, and distressed woman); positive pictures (valence: *M* = 6.98, *SD* = 1.61; arousal: *M* = 4.35, *SD* = 2.17; e.g., laughing boy, grateful girl, happy elderly woman, and pilot); and neutral pictures (valence: *M* = 5.27, *SD* = 1.45; arousal: *M* = 3.57, *SD* = 1.96; e.g., neutral faces of woman, man, and child). The full list of pictures for each valence arousal ratings are presented in the Supplemental Material. There were equal numbers of male and female faces in each of the three experimental conditions. Stimuli were presented in color and standardized in size to 507 x 635 pixels, against a gray background.

The task was presented in three runs. Each run included three blocks (cognitive empathy, affective empathy, and age perception control) of positive, negative, and neutral images, resulting in nine blocks in total per run. The order of conditions in each run was pseudo-randomized. Positive, negative, and neutral stimuli were counterbalanced across experimental conditions and runs (i.e., positive pictures presented in the affective empathy condition in Run 1 were presented in the cognitive and control conditions in Run 2 and 3, respectively). We ensured that each block of positive, negative, or neutral pictures was only presented once within each run. The order of presenting each run was counterbalanced across participants. To enhance design efficiency (Huettel et al., 2014), each run included two low-level blocks, presented randomly for 42 seconds during the run, consisting of a fixation cross on gray background. In addition, a jittered fixation cross was presented between each block in each run randomly from one of the following durations: 1.5, 2, and 2.5 seconds. Each run lasted 7.7 minutes, for a total task duration of 23.1 minutes. We used Psychtoolbox for task programming, presentation, and response recording.

**Figure 1.**
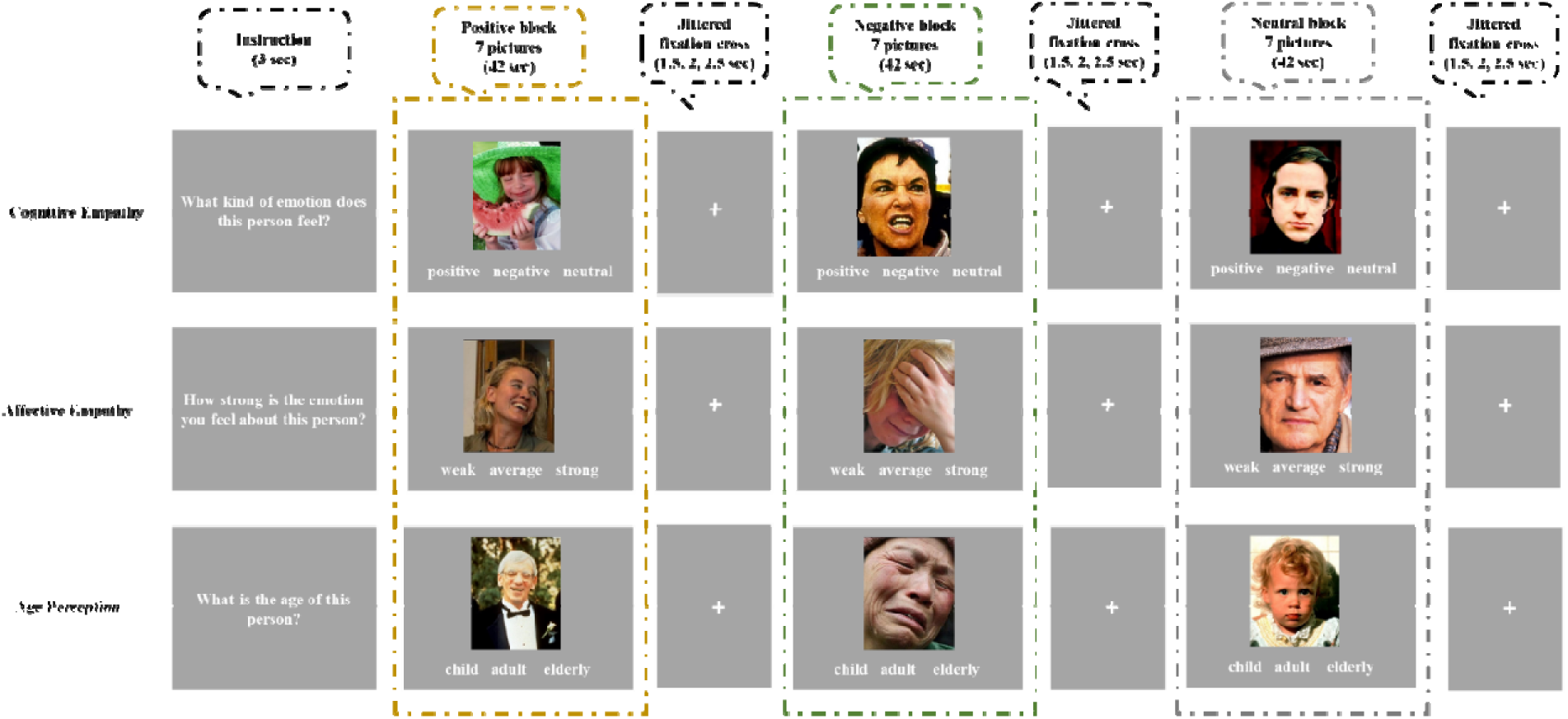
Experimental task. The Multifaceted Empathy Test (MET) included three experimental conditions: cognitive empathy, affective empathy, and age perception control. Each condition consisted of three blocks of either positive, negative, or neutral images. Each condition started with a 3-second instruction, followed by a presentation of 7 images, each for 6 seconds, for a total of 42 seconds per block. Following each block, a jittered fixation cross (duration: 1.5, 2, or 2.5 seconds) was presented.

### Background measures

In the behavioral/neuropsychological test session, participants completed a series of tasks pertaining to executive functioning: the Stroop Task (Jensen & Rohwer, 1966), the abbreviated version of the Raven’s Progressive Matrices (Bilker et al., 2012), the Trail Making Test (Reitan & Wolfson, 1986), and a verbal fluency measure (Newcombe, 1969). Emotional well-being was measured with the Depression, Anxiety, Stress Scale – DASS-21 (Lovibond & Lovibond, 1995). In addition, the Empathy Quotient (Baron-Cohen & Wheelwright, 2004), the Interpersonal Reactivity Index (Davis, 1983), and the Reading the Mind in the Eyes test (Baron Cohen et al. (2001) measured empathy and theory of mind. As shown in Table 1, there were no differences between the two age groups in background measures, with the exception of the three DASS-21 subscales, for which younger adults reported higher levels than older adults, and the personal distress subscale from the IRI, for which younger adults scored higher than older adults (*ps* < 0.001).^1^

### MRI image acquisition

Functional images were acquired at the Centre for Advanced Imaging on a 3T Siemens Prisma scanner with a 32-channel head coil, using a whole-brain T2*-weighted multiband sequence (834 interleaved slices, repetition time (TR) = 612 ms, echo time (TE) = 30 ms, voxel size = 2.5 mm^3^, field of view (FOV) = 190 mm, flip angle = 52°, multi-band acceleration factor = 5). High-resolution T1-weighted images were acquired with an MP2RAGE sequence (176 slices per slab, TR = 4000 ms, TE = 2.91 ms, TI = 700 ms, voxel size = 1 mm^3^, FOV = 256 mm, PAT mode = GRAPPA). Also, a diffusion-weighted imaging sequence with two shells was conducted (shell one: TR = 4100 ms, TE = 70 ms, voxel size = 2 mm^3^, number of slices =68, FoV = 244 mm, b-value: 2500 s/mm^2^, 66 directions; shell two: TR = 4100 ms, TE = 70 ms, voxel size = 2 mm^3^, number of slices =68, FoV = 244 mm, b-value: 1200 s/mm^2^, 33 directions). To minimize noise and head movement, participants were provided with earplugs and cushions around their head inside the head coil. Participants were presented with the task on a computer screen through a mirror mounted on top of the head coil.

## Data analysis

### Behavioral data

We conducted a repeated measures analysis of variance (ANOVA) on mean response times with experimental condition (cognitive empathy, affective empathy, age perception control) as well as valence (positive, negative, neutral) as within-subject factors and age group (younger, older) as between-subject factor. While both response times and accuracy were collected during the task, given that the block design presented blocks of seven positive, negative, or neutral images respectively, accuracy did not vary within a block and thus was limited as an outcome variable^2^. Additionally, while accurate responses were possible during the cognitive empathy and the age perception control conditions, during the affective empathy condition participants were instructed to indicate how strongly they felt towards the stimuli and thus were not probed on accuracy. Therefore, reaction times were used to reflect behavioral performance in the MET. Response times were skewed as determined by the Shapiro-Wilk normality test and thus we used log transformed response times in the analyses reported here (see Supplemental Material for details on normality test results and distributions).

We additionally investigated relationships between the background measures with the structural and functional measures. The results of these analyses are reported in the Supplemental Material.

### fMRI

*Preprocessing.* T2*-weighted images were preprocessed with Statistical Parametric Mapping Software (SPM12; http://www.fil.ion.ucl.ac.uk/spm) implemented in MATLAB 2017 (Mathworks Inc., MA). Following realignment to a mean image for head-motion correction, images were segmented into gray and white matter. Images were spatially normalized into a standard stereotaxic space with a 2-mm^3^ voxel size, using the Montreal Neurological Institute (MNI) template and spatially smoothed with a 6-mm^3^ Gaussian Kernel.

*Analyses.* We used task Partial Least Squares (PLS; McIntosh et al., 1996; McIntosh et al., 2004), as implemented in the PLS software running on MATLAB 2017 (The MathWorks Inc., MA), to determine age-related differences in whole-brain activity patterns for the three experimental conditions (cognitive empathy, affective empathy, age perception) and by image valence (positive, negative, neutral).

PLS is a model-free, multivariate analytical technique (for a detailed tutorial and review of PLS, see Krishnan et al. (2011)) that allows examination of the relationship between brain activity and multiple experimental conditions simultaneously and that does not require multiple comparison correction (McIntosh et al., 2004). For the whole-brain analysis, we included all three experimental conditions: cognitive empathy, affective empathy, and age perception for both age groups. PLS captures the pattern of covariance between data that are unique to the data set without imposing arbitrary contrasts for or assumptions about the experimental conditions. PLS organizes all data from all participants and all experimental conditions into a single matrix, and by using a singular value decomposition (SVD) finds a set of orthogonal latent variables (LVs), which represent linear combinations of the original variables.

In contrast to the more commonly used General Linear Model, PLS not only considers the temporal relationship between the fMRI data and the task design but also the spatial relationship between activated voxels. As a spatio-temporal analysis method, PLS is based on the joint variance of individual voxels and is, thus, more sensitive to the covariance of the brain activity. By using PLS, our results here are not based on contrasts that show regions that are more or less engaged during one condition over another (i.e., our results do not follow the logic of the subtraction method). Rather, our results reflect changes in brain activity related to task manipulation uncovering the brain’s responses to differences between the experimental conditions. Thus, our study design leverages the strengths of PLS for a novel multivariate examination of age-related differences in cognitive and affective empathic response to positive and negative emotions.

Each LV identified with PLS delineates brain activity patterns related to the experimental conditions. Usually, the first LV accounts for the largest covariance of the data, with a progressively smaller amount of covariance for subsequent LVs. The amount of covariance accounted by an LV is referred to as singular value. Each LV consists of a spatio-temporal pattern of brain activity (referred to as voxel saliences), a set of weights that indicates the relationship between the brain activity and the experimental conditions (referred to as task saliences). Each LV contains brain scores that reflect how each participant contributed to the pattern expressed in the respective LV. A permutation test with 500 random reordering and resampling was conducted to infer the statistical significance of each LV (McIntosh et al., 1996). Additionally, the robustness of voxel saliences was assessed with a bootstrap estimation with 100 resampling iterations^3^ (Efron & Tibshirani, 1985). Peak voxels with a bootstrap ratio (i.e., salience/standard error) ≥ 2.5 were considered reliable, as this approximates *p* < 0.005 (Sampson et al., 1989).

In this study, we used a block design analysis by defining, within each of the three experimental conditions (cognitive empathy, affective empathy, age perception), the onset of each block of positive, negative, and neutral stimuli, respectively. For the *Hypothesis 1* set, to determine age-related differences in brain activity patterns for cognitive and affective empathy, we included both age groups and all three experimental conditions, irrespective of valence. For the *Hypothesis 2* set, to examine the role of valence within each experimental condition, we conducted separate analyses for each of the three experimental conditions with positive, negative, and neutral images and both age groups included in each analysis. To determine associations between structural integrity and functional network activity, we examined whether fractional anisotropy (FA values) of anterior and posterior cingulum bundle white matter tracts were correlated with functional networks activated during the MET and whether this association varied by stimulus valence and age group.

### DTI

The *recon-all* command implemented in FreeSurfer (v6.0) (http://surfer.nmr.mgh.harvard.edu/) was used for the segmentation of T1-weighted images (Dale et al., 1999). The diffusion-weighted (DW) data were preprocessed to correct for head movements, eddy current distortions, and signal intensity inhomogeneities, using tools implemented in MRtrix3 (Tournier et al., 2012). DW and T1-weighted images were co-registered using boundary-based registration (Greve & Fischl, 2009). A five-tissue-type segmented image (cortical grey matter, white matter, sub-cortical grey matter, cerebrospinal fluid, pathological tissue) was generated from the preprocessed T1-weighted images. Response functions were estimated using a multi-shell, multi-tissue algorithm, and multi-tissue constrained spherical deconvolution was applied to obtain fiber orientation distributions (FOD) (Jeurissen et al., 2014). For the reconstruction of the anterior and posterior cingulum subdivisions, we used a deterministic tractography algorithm based on spherical deconvolution, which takes the FOD image as input and samples it at each streamline step. Newton optimization was performed on the sphere from the current streamline tangent orientation to locate the orientation of the nearest FOD amplitude peak. The step size of the tracking algorithm was set to 0.5mm, with a cut-off value for the FOD amplitude of 0.05, maximum turning angle of 45°, and minimum path length of 10mm. Mean FA was calculated for each reconstructed tract, as a general marker of integrity within white matter structures, suggesting coherence within a fiber and voxel density (Beaulieu, 2002).

### Tractography pipeline

Anatomical landmarks were identified on color-coded diffusion tensor maps. An exclusion region of interest (ROI) was drawn across the mid-sagittal plane to exclude interhemispheric projections. Further exclusion ROIs were drawn to exclude tracts that deviated from the anatomy of the cingulum bundle. All tracts were reconstructed in the left and right hemisphere. The anterior and posterior subdivisions were reconstructed as described previously (Metzler-Baddeley et al., 2012) with minor modifications: The anterior cingulum was defined as the cingulum segment rostral to the anterior commissure. The seed ROI was drawn in line with the anterior commissure in the coronal plane. One inclusion ROI was placed in the slice where the most inferior part of the genu was identified in the axial plane; another inclusion ROI was drawn in the coronal plane where the most posterior part of the genu was visible. The posterior cingulum was defined as the cingulum segment caudal to the posterior commissure. The seed ROI was placed in line with the posterior commissure in the coronal plane. One inclusion ROI was drawn in the slice where the most inferior part of the splenium was identified in the axial plane; another inclusion ROI was placed in the coronal plane where the most anterior part of the splenium was visible.

### Structure-function analysis

The structure-function analysis followed previous approaches (Ziaei et al., 2020). Figure 2 outlines our analysis procedure. In particular, to determine age-differential associations between structural integrity (FA values) of the cingulum bundle tracts and whole-brain activation during the MET, we performed separate PLS analyses for each white matter structure (i.e., anterior, posterior subdivision of the cingulum tract). Thus, the results reflect respective correlations between brain scores and FA values of the *(i)* anterior cingulum, the *(ii)* posterior cingulum, separately for the left and right hemisphere. In particular, the respective FA values for each participant were correlated with the respective brain scores for each participant in the cognitive and affective empathy conditions (in separate analyses), including positive, negative, and neutral images in the models. The correlations were acquired for each participant within each age group. These analyses aimed to examine the relationship between functional brain activity pattern and white matter tract FA values in both younger and older adults.

**Figure 2.**
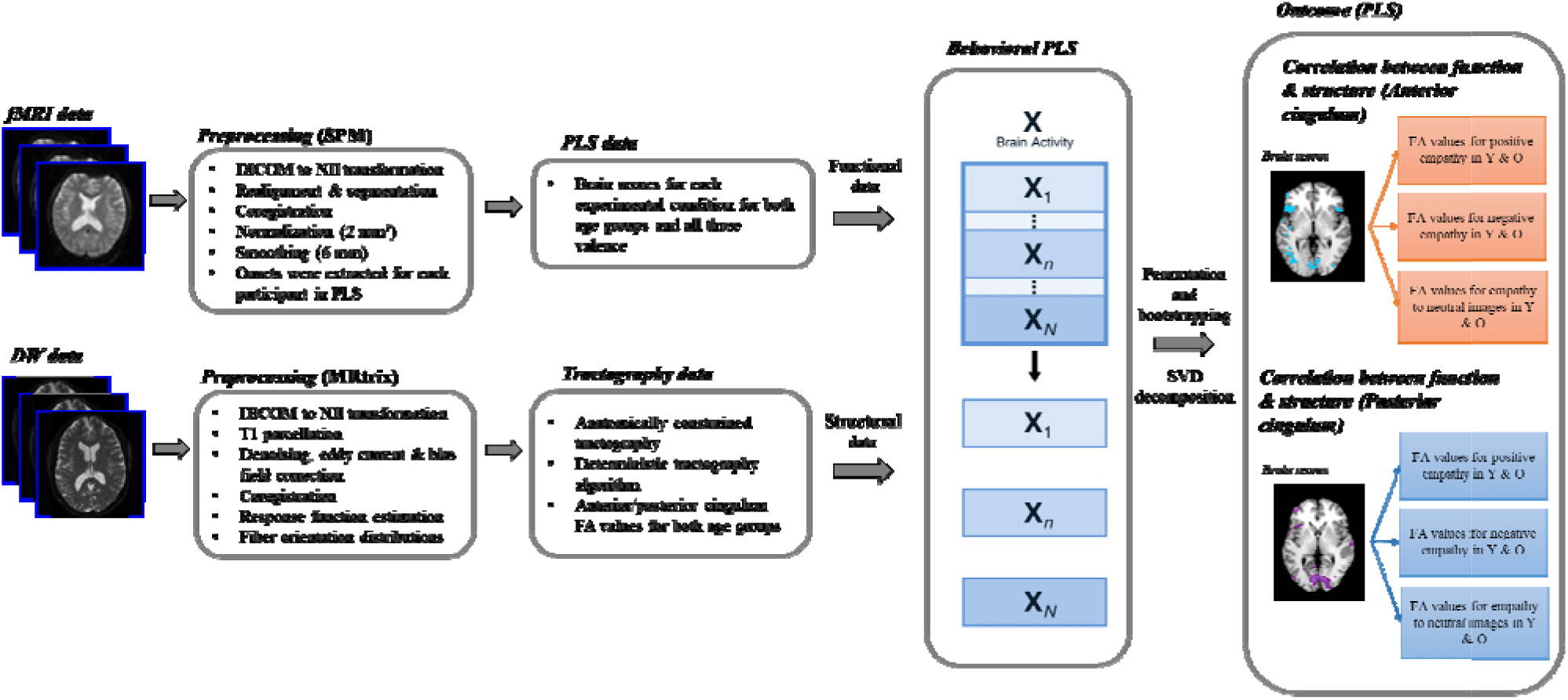
Methodological approach. Following preprocessing of fMRI and DTI data, brain scores and cingulum fractional anisotropy (FA) values were entered into the model to delineate latent variables that best described the relationship between structural and functional data. fMRI = Functional Magnetic Resonance Imaging; DW = Diffusion Weighted; SPM = Statistical Parametric Mapping; DICOM = Digital Imaging and Communications in Medicine; NII = NIFTI; PLS = Partial Least Square; FA = Fractional Anisotropy; AC = Anterior Cingulate; PC = Posterior Cingulate; Y = Younger Adults; O = Older Adults.

## Results

### Behavioral performance

#### Response times

All three main effects were significant. The main effect for experimental condition (*F*(2,96) = 89.54, *p* < 0.001, 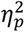 = 0.65) showed that participants overall responded slower in the affective (*M* = 2.20, *SD* = 0.54) than the cognitive (*M* = 1.93, *SD* = 0.41) condition. The main effect for valence (*F*(2,96) = 40.95, *p* < 0.001, 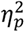 = 0.46) showed that participants overall responded faster to positive (*M* = 1.70, *SD* = 0.36) than negative (*M* = 1.87, *SD* = 0.35) stimuli. The main effect for age group (*F*(1,48) = 13.15, *p* = 0.003, 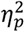 = 0.21) revealed that overall older adults (*M* = 2.03, *SD* = 0.32) responded slower than younger adults (*M* = 1.71, *SD* = 0.29) (Figure 3).

A significant experimental condition by age group interaction (*F*(2,96) = 8.36, *p* < 0.001, 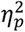 = 0.14) furthermore indicated that older compared to younger adults responded slower during affective (*t*(49) = 2.61, *p* = 0.012, *d* = 0.74), and particularly during cognitive (*t*(48) = 5.33, *p* < 0.001, *d* = 1.53), empathy, while the age groups did not differ in response time during the age perception control condition (*t*(49) = 0.95, *p* = 0.34, *d* = 0.27). Also, a significant valence by age group interaction (*F*(2,96) = 4.34, *p* = 0.016, 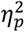 = 0.08) showed that relative to younger participants, older adults responded slower to negative than positive stimuli (*t_older adults_* (23) = 4.58, *p* < 0.001, *d* = 1.9). A significant experimental condition by valence interaction (*F*(4,192) = 36.53, *p* < 0.001, 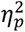 = 0.43), finally, showed that participants responded slower to neutral than emotional stimuli during cognitive empathy (*t*(49) = 9.66, *p* < 0.001, *d* = 2.76), while response time was not different during either the affective empathy or the age perception control condition (all *p*s > 0.05).

**Figure 3.**
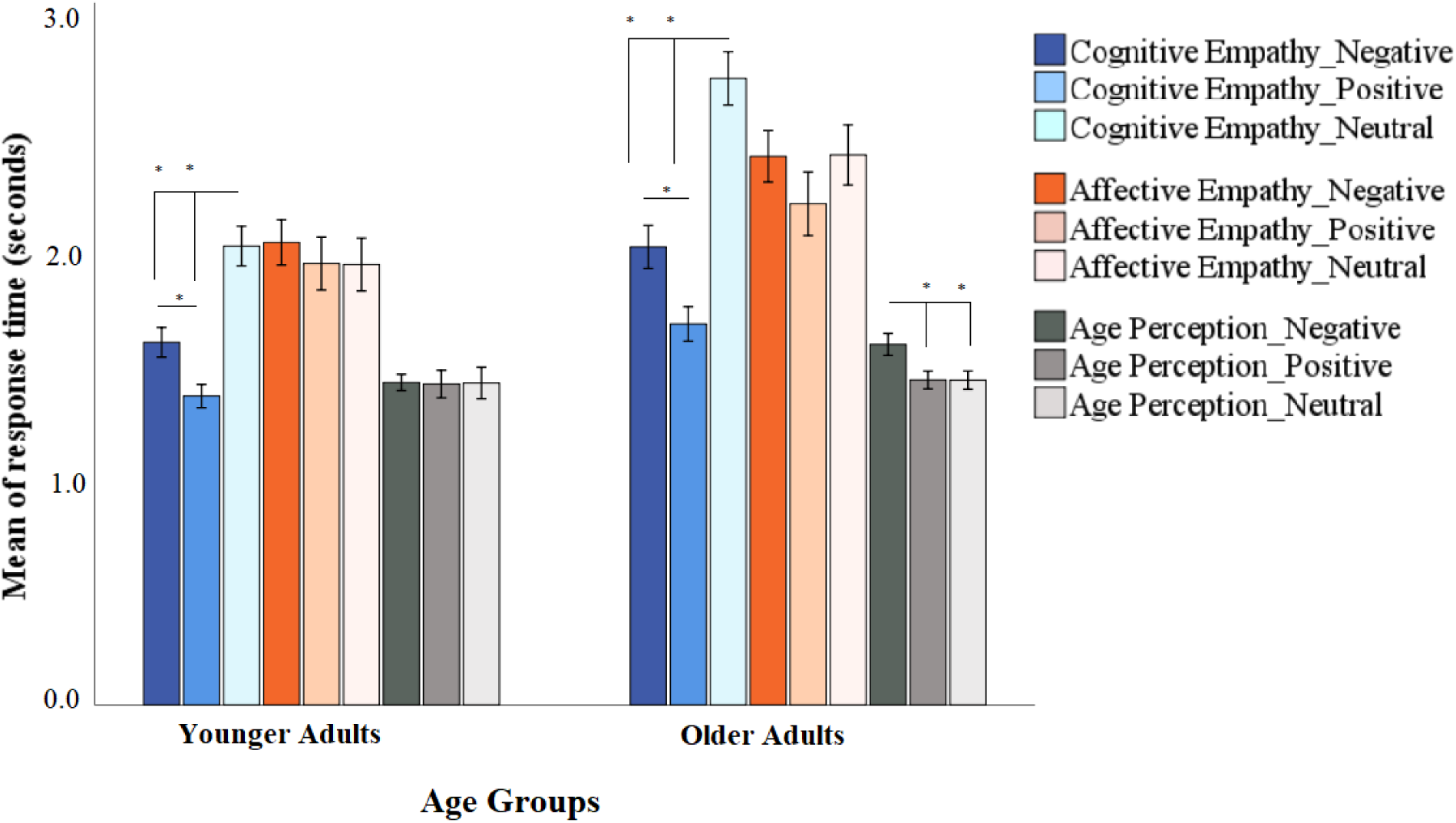
Reaction times across all conditions and age groups. Panel A. Mean response times (in seconds) for the three experimental conditions (cognitive empathy, affective empathy, age perception control) in younger and older adults for positive, negative, and neutral stimuli. N = number; Error bars = +/-1 standard error. * denotes *p* < 0.000.

### Whole-brain activity pattern

#### Age-related differences in cognitive and affective empathy (Hypotheses 1a&b)

This analysis resulted in two significant LVs. The first LV accounted for 48% of the covariance in the data (*p* < 0.001). This brain pattern included the bilateral insula, bilateral parahippocampus, right superior frontal gyrus, left superior temporal gyrus, right anterior cingulate, left medial frontal gyrus, bilateral precuneus, and bilateral posterior cingulate (Figure 4A). This network was engaged more by older than younger participants (Figure 4B).

The second LV accounted for 26% of the covariance in the data (*p* < 0.001). This mainly left-sided brain pattern included the left inferior frontal gyrus, left superior gyrus, left medial frontal gyrus, left inferior parietal lobe, bilateral anterior cingulate, left supramarginal gyrus, left superior temporal gyrus, left middle temporal gyrus, and bilateral insula (Figure 4C). Both age groups engaged this network similarly during the affective empathy condition (Figure 4D). Additionally, this LV included another pattern, which corresponded to the age perception control condition similarly in both age groups and included the right pre and post central gyrus, bilateral inferior parietal lobe, and right precuneus.

**Figure 4.**
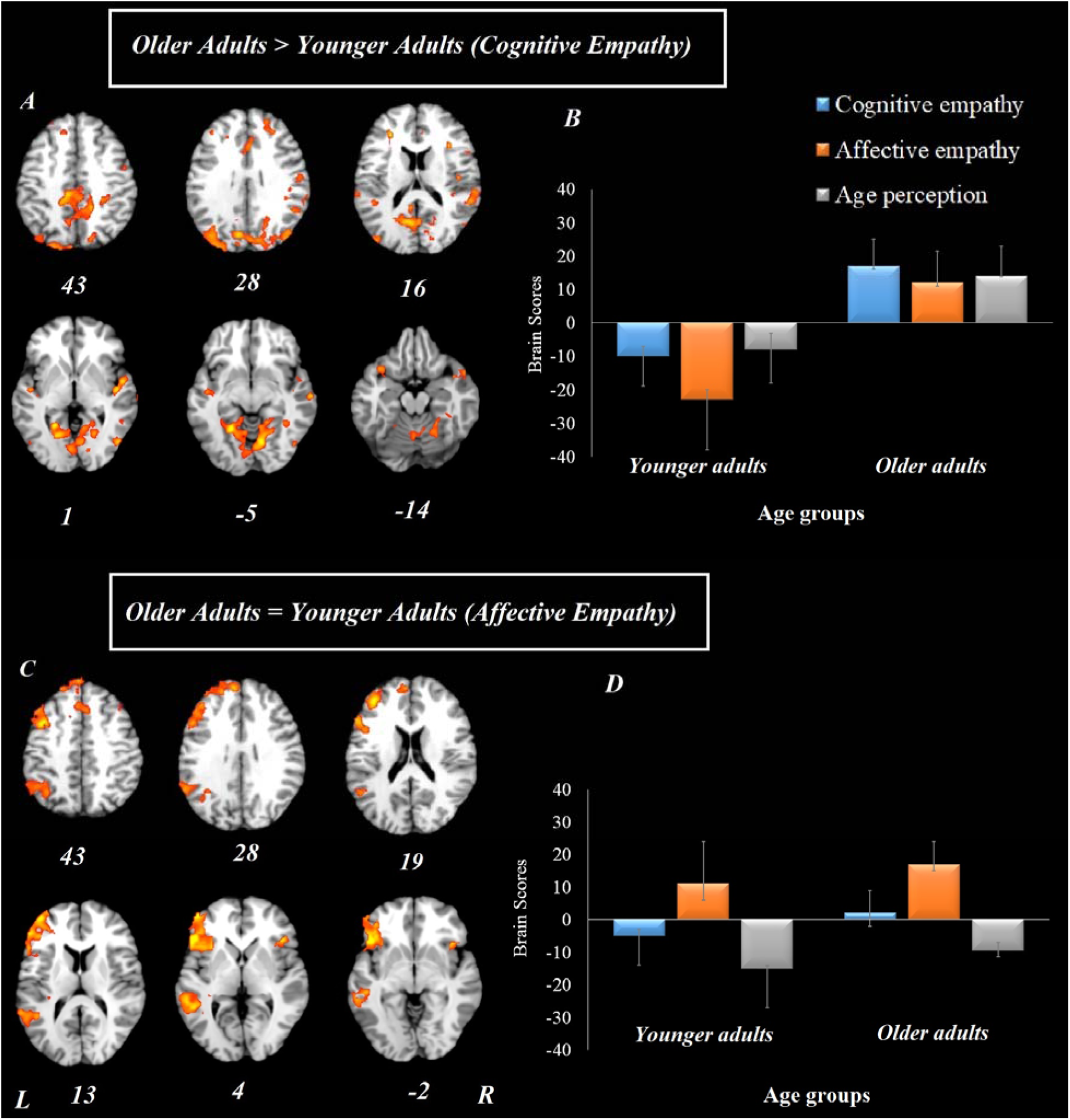
Results from the whole-brain analysis for the three experimental conditions (cognitive empathy, affective empathy, age perception control) in younger and older adults. Panel A represents the brain activity pattern across all three experimental conditions, differentiating between younger and older adults. Panel B represents brain scores associated with the brain activity pattern presented in Panel A. Panel C represents the brain activity pattern differentiating affective empathy from the other two experimental conditions, similarly for younger and older adults. Panel D represents brain scores associated with the brain activity pattern presented in Panel C. Error bars represent confidence intervals at 95%. For all reported regions a bootstrap ratio of ≥ 2.5 and cluster size of ≥ 50 voxels was applied. L = left hemisphere, R = right hemisphere.

#### Age-related differences in valence effects on cognitive and affective empathy (Hypotheses 2a&b)

The analysis pertaining to cognitive empathy resulted in one significant LV that accounted for 39% of the covariance in the data (*p* = 0.002). This network included the bilateral inferior frontal gyrus, anterior cingulate, bilateral superior temporal gyrus, bilateral superior frontal gyrus, medial frontal gyrus, and bilateral precentral gyrus regions (Figure 5A). This network was positively correlated with cognitive empathy to negative and neutral stimuli only in older adults (Figure 5B).

**Figure 5.**
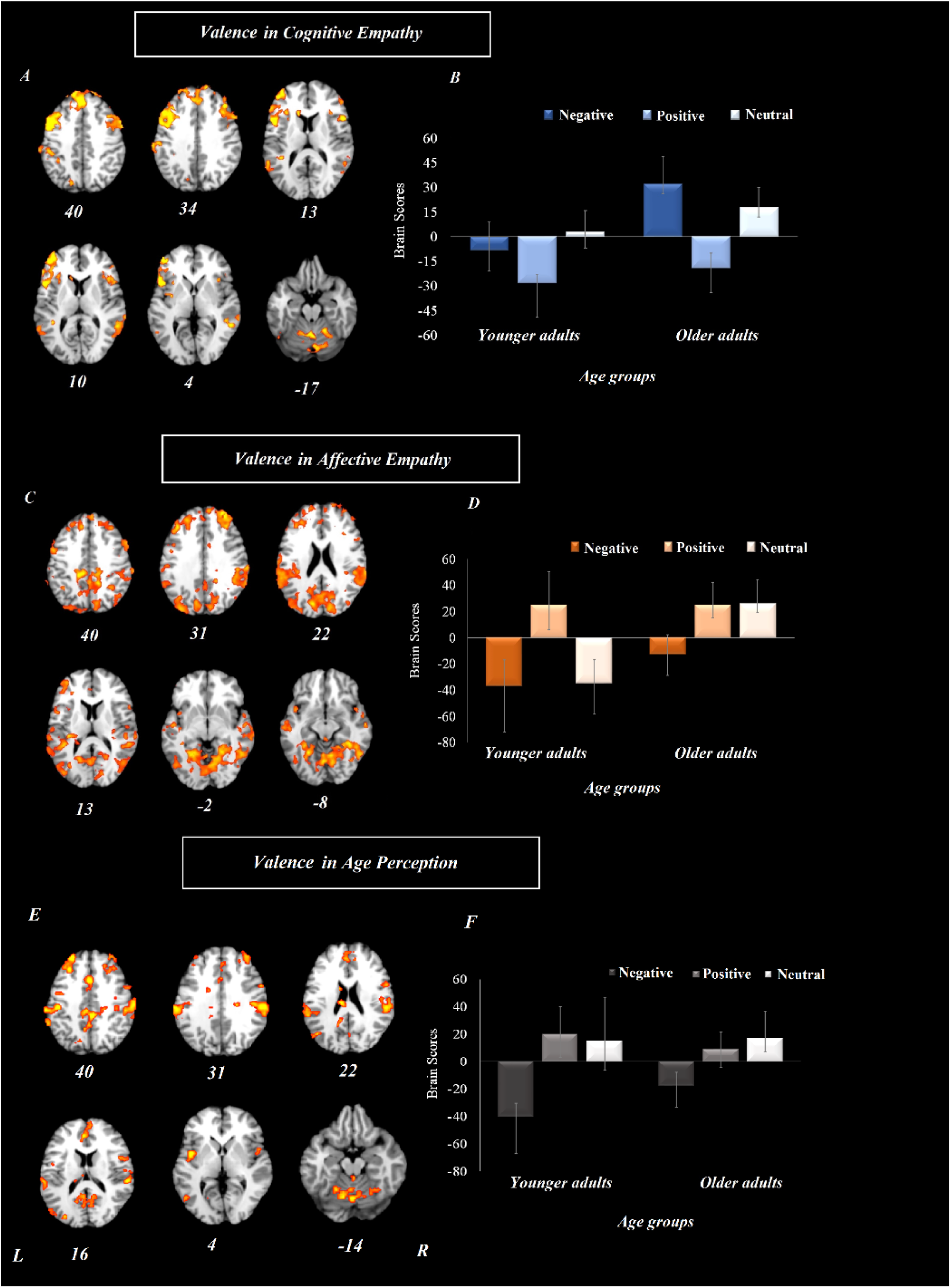
Results from the whole-brain analysis regarding valence modulation (positive, negative, neutral) for cognitive empathy, affective empathy, and age perception control in younger and older adults. Panel A refers to cognitive empathy and represents the brain activity pattern differentiating negative and neutral from positive stimuli in older adults. Panel B represents brain scores associated with the brain activity pattern presented in Panel A (cognitive empathy). Panel C refers to affective empathy and represents the brain activity pattern differentiating positive stimuli from negative and neutral in younger adults and positive and neutral from negative stimuli in older adults. Panel D represents brain scores associated with the brain activity pattern presented in Panel C (affective empathy). Panel E refers to age perception and represents the brain activity pattern differentiating negative from positive and neutral stimuli in younger adults and neutral from negative stimuli in older adults. Panel F represents brain scores associated with the brain activity pattern presented in Panel E (age perception). Error bars represent confidence intervals at 95%. For all reported regions a bootstrap ratio of ≥ 2.5 and cluster size of ≥ 50 voxels was applied. L = left hemisphere, R = right hemisphere.

The analysis pertaining to affective empathy also resulted in one significant LV which accounted for 51% of the covariance in the data (*p* < 0.001). This network included the bilateral middle and superior frontal gyrus, posterior cingulate, precuneus, lingual gyrus, inferior parietal lobe, and superior temporal gyrus (Figure 5C). This wide-spread network was positively correlated with affective empathy to positive stimuli in both younger and older adults, and additionally to neutral stimuli in older adults (Figure 5D).

The analysis pertaining to the age perception control condition resulted in one significant LV that accounted for 45% of the covariance in the data (*p* = 0.012). This network included the anterior and posterior cingulate cortex, right inferior frontal gyrus, bilateral middle frontal gyrus, bilateral inferior parietal lobe, bilateral insula, bilateral superior temporal gyrus, and right precuneus (Figure 5E). This network was positively correlated with age perception of positive stimuli in younger adults and neutral stimuli in older adults (Figure 5F).

#### Structure-function relationship

We next examined the relationship between anterior cingulum bundle and posterior cingulum bundle microstructure and brain activity during the MET, by valence and age group.

##### Anterior cingulum bundle

Our analysis testing associations between anterior cingulum bundle FA values and brain activation during *cognitive empathy* revealed one significant LV which accounted for 25% of the covariance in the data (*p* = 0.008). This network included the left superior frontal gyrus, bilateral superior temporal gyrus, posterior and anterior cingulate. Younger adults with higher FA values in the left anterior cingulum bundle recruited this functional network in cognitive empathic responding to negative stimuli.

Additionally, our analysis testing associations between anterior cingulum bundle FA values and brain activation for *affective empathy* revealed one significant LV which accounted for 25% of the covariance in the data (*p* < 0.001). This network included the left superior frontal gyrus, bilateral insula, left superior temporal gyrus, bilateral parietal lobe, left hippocampus, left caudate, and left fusiform gyrus. Older adults with higher FA values in the right anterior cingulum bundle recruited this functional network in affective empathic responding across all three emotional stimuli. No other effects were reliable (all confidence intervals crossed zero; supplementary Figure 3).

##### Posterior cingulum bundle

Our analysis testing associations between posterior cingulum FA values and brain activity during *cognitive empathy* revealed one LV which accounted for 27% of the covariance in the data (*p* = 0.008). This network included bilateral middle frontal gyrus, superior frontal gyrus, anterior cingulate, bilateral inferior parietal lobe, and right superior temporal gyrus. Younger adults with higher FA values in the left posterior cingulum bundle recruited this functional network in cognitive empathic responding to negative stimuli.

Additionally, as shown in Figure 6, our analysis testing associations between posterior cingulum bundle FA values and brain activation for *affective empathy* revealed one significant LV which accounted for 28% of the covariance in the data (*p* = 0.004). This network included the bilateral ventromedial prefrontal cortex, medial prefrontal cortex, and bilateral cuneus (Figure 6B). Older adults with higher FA values in both left and right posterior cingulum bundle recruited this functional network in affective empathic responding to positive stimuli (top and bottom panels Figure 6C). In younger adults this structure-function relationship was only reliable for the right (bottom panel Figure 6C) but not left (top panel Figure 6C) posterior cingulum tract.

**Figure 6.**
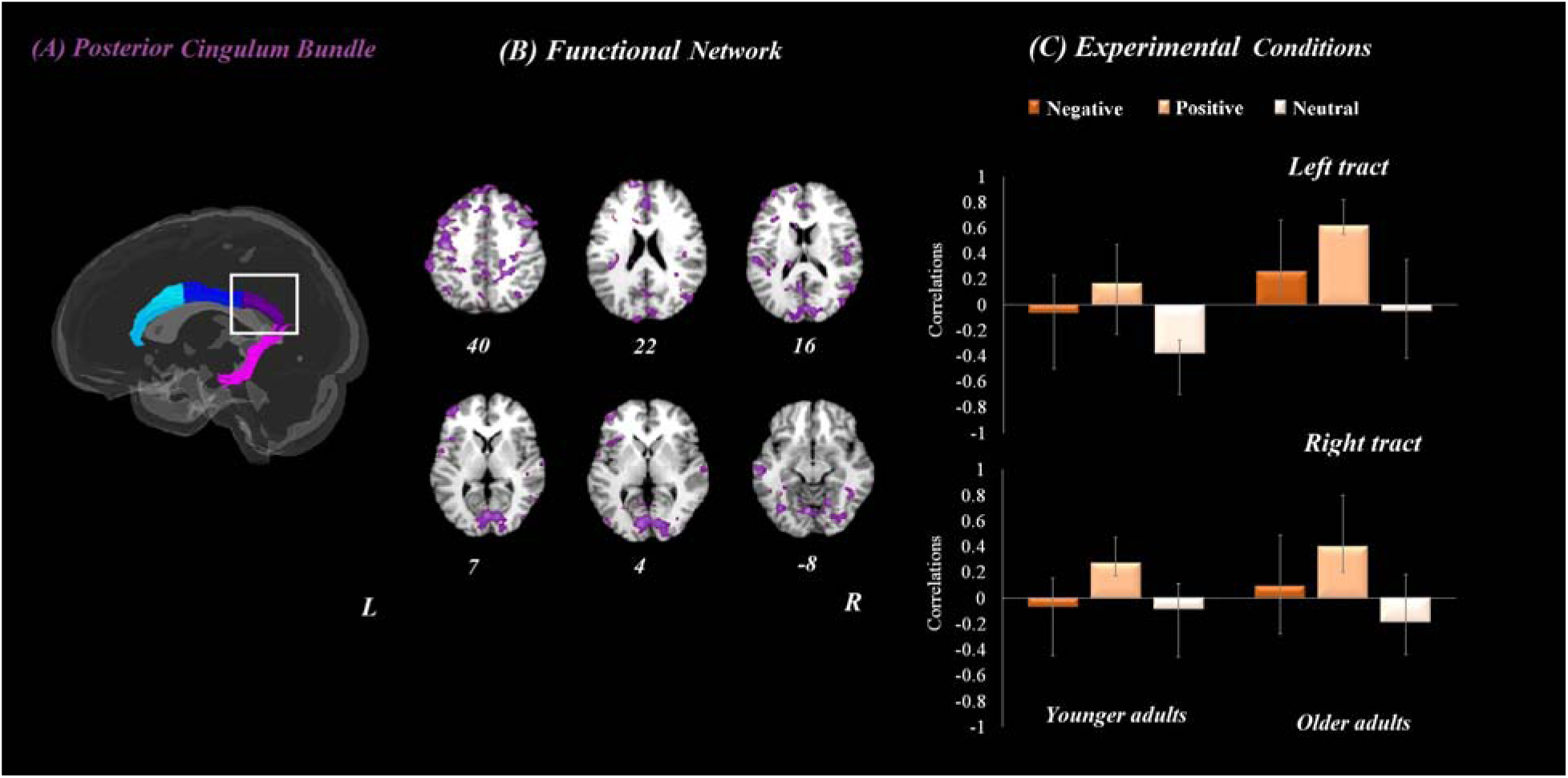
Results from whole-brain analyses and posterior cingulum fractional anisotropy (FA) values in both age groups for the affective empathy condition. Panel A depicts the posterior cingulum bundle tract. Panel B refers to the brain activation pattern which was correlated with the posterior cingulum bundle FA values during affective empathy. Panel C refers to the correlation between posterior cingulum bundle FA values and brain patterns for the three valence conditions in both younger and older adults. Error bars represent confidence intervals at 95%. For all reported regions a bootstrap ratio of ≥ 2.5 and cluster size of ≥ 50 voxels was applied. L = left hemisphere, R = right hemisphere.

## Discussion

Age-related differences in empathy have been researched across multiple studies, and overall a pattern has emerged suggesting decrease in cognitive empathy with age, while effects on affective empathy are more mixed (see also Bailey et al., 2021; Ebner et al., 2017, for overviews). Going beyond previous research by examining not only behavior but also brain structure and function in cognitive and affective empathy among younger and older adults, this study generated several novel insights.

Behaviorally, we found that older adults’ empathic responding was slower than their responses to the age perception control condition; and their response times were affected by the valence of the stimulus during cognitive, but not affective, empathy. At the neural level, we did not find any evidence for age-related reduced activity during cognitive empathy, thus not supporting *Hypothesis 1a*. Supporting *Hypothesis 1b*, however, we found that for affective empathy both age groups recruited a similar brain network. We also found that older, but not younger, adults engaged regions of the salience network in response to negative emotions during cognitive empathy, supporting *Hypothesis 2a*. In contrast, with regards to affective empathy, both age groups, and not only older adults, engaged a similar pattern of brain regions that contained nodes of the default mode network in response to positive emotions, partially supporting *Hypothesis 2b*. Our structure-function analyses, finally, revealed that microstructure of the posterior, but not the anterior, cingulum bundle was related to engagement of major nodes of the default mode network during affective empathy to positive stimuli in both age groups. These central findings from this work will be discussed in more detail next.

The results of our whole-brain analysis (the first LV) indicated that regions such as bilateral insula, right superior frontal gyrus, left superior temporal gyrus, right anterior cingulate, left medial frontal gyrus, bilateral precuneus, and bilateral posterior cingulate were engaged across all experimental conditions among older adults, perhaps reflecting general cognitive effort needed for all conditions. Some of these regions have been implicated in empathic responding (e.g., insula and anterior cingulate; Bellucci et al., 2020) and understanding others’ mental states (e.g., superior temporal gyrus; Gallagher & Frith, 2003; et al., 2020). In addition, however, they have also been involved in exerting cognitive effort during a task (e.g., superior frontal gyrus; Aron et al., 2014) as well as in self-referential processing (possibly in relation to others) more generally (e.g., precuneus and posterior cingulate; Buckner et al., 2008), in support of the notion that these areas reflect general domain processing rather than experimental condition specific processing.

Neither our brain nor our behavioral data supported age-related differences in affective empathy. Thus, in contrast to Chen et al. (2014), we did not observe reduced engagement of core empathy regions including the anterior insula, mid-cingulate, and medial prefrontal cortex in older adults. Rather, both younger and older adults in our study recruited these regions similarly during the affective empathy task. Differences in the type of stimuli used in our study (i.e., non-pain stimuli) compared to the pain stimuli used in Chen et al. (2014) may underlie these divergent findings. Speaking against this methodological explanation, a recent meta-analysis found similarities in empathic responses to pain and non-pain stimuli (Timmers et al., 2018); this comparability across stimulus types, however, has not been confirmed in research with older individuals yet. Additionally, it has to be noted that Chen et al. (2014) investigated empathic responses in three age groups, younger, middle age, and older adults; and while the neural correlates of each age group were compared in their study, the use of different stimuli and a sample from quite a different population than the present study’s sample has to be noted when comparing Chen et al. (2014)’s results with the results from the current study. Further, they conducted univariate, contrast-based imaging analyses, while the present study adopted a multivariate analysis method.

Our functional MRI findings for affective empathy do not support the notion of diminished internal bodily perception (interoceptive awareness) among older adults (Mendes, 2010), but functional sparing for affective processing in aging, as observed here, is in line with the brain maintenance hypothesis discussed in the cognitive aging literature (Nyberg et al., 2012). One could also argue that the idea of functional reserve that protects older adults from decline does especially well apply as an explanation to our findings given that participants in this study were cognitively high functioning (see background measures in Table 1). Thus, they may have had high cognitive reserve, which could have resulted in performance on a level comparable with the younger participants – and especially so for the affective empathy task which may have been easier to perform than the cognitive empathy task. Future research will be able to test this interpretation.

We observed age-related differences in valence modulation during cognitive empathy, both on the behavioral and the neural level. In particular, we found that older adults recruited a neural network that comprised bilateral insula and anterior cingulate, core nodes of the salience/midcingulo-insular network network (Menon, 2015; Menon & Uddin, 2010), more in their cognitive empathic response to negative stimuli than younger adults did. This finding aligns with behavioral evidence that older compared to younger adults experience greater difficulty processing negative than positive emotions (Hayes et al., 2020; Ruffman et al., 2008). In line with evidence from these meta-analyses, processing of negative (relative to positive) emotions may be more effortful and considered more threatening for older adults, as it does not align with their primary goals and implicit motivations. As a result, enhanced prefrontal cortex and insula activation for processing negative relative to positive information among older adults (Ebner et al., 2012; Ziaei et al., 2016) may be reflective of greater cognitive control and/or emotional down-regulatory processes at work.

Another possibility for engagement of the salience network during cognitive empathy to negative emotions in older adults is that this network, and insula specifically, is involved in orienting attention towards relevant stimuli in the environment (Menon & Uddin, 2010). Given the salience of negative emotions and their importance for survival, orienting attention towards negative emotions is crucial and, therefore, associated with insular response. Moreover, insular activity in this context may be reflective of a brain response that is commonly observed across various cognitive tasks to guide behavior in dynamic social contexts where recognition of negative emotions is crucial (Bernhardt & Singer, 2012). Our result, however, contradicts suggestions of age-related reductions in insular activation subserving interoception and the simulation of emotions in others (Mather, 2016). Certainly, further investigation is needed to determine the relationship between bodily response (such as heart rate variability and skin conductance) and insular activity during empathy, and social cognition more broadly, across adult development and in aging.

Our finding of no age-related difference in affective responding to positive stimuli is in line with socioemotional selectivity theory (SST) (Carstensen et al., 2003; Carstensen et al., 1999), which proposes that older adults preferentially process positive over negative stimuli. In other words, older adults’ bias towards positive emotions may have facilitated processing of positive emotions in the affective empathy task, resulting in comparable brain and behavioral activity patterns between the age groups under this condition. Thus, the present study’s results could be interpreted as suggesting that the positivity effect reported in the aging literature for various cognitive and social-cognitive processes also extends to affective empathy. In fact, this finding also aligns well with the motivational empathy account (Weisz & Zaki, 2018). That is, older adults might be more motivated to process positive than negative emotions, as these emotions correspond with their social goals.

Another explanation for similar responses among the age groups towards positive affective empathy might be a general tendency in humans, at any age, to show empathic responses towards positive emotions. Relatedly, considering evidence that it is easier to show support for a partner’s positive than negative life events (Andreychik, 2019; Gable et al., 2006), it is possible that the age groups do not differ in the neural network involved under this condition. Recent neuroimaging evidence furthermore supports the idea that positive empathy makes prosocial acts feel more rewarding, activating regions such as the ventromedial prefrontal cortex (Harbaugh et al., 2007; Hare et al., 2010). Thus, empathizing with positive emotions may be more rewarding and/or easier, generally, and comparably so for younger and older adults.

Taken together, the findings reported here well align with previous studies supporting salience network/ midcingulo-insular network (including the insula and the cingulate cortex) activation during cognitive empathy (Bernhardt & Singer, 2012; Pasquini et al., 2020; Rankin et al., 2006). Positive empathy can engage areas related to the processing of positive emotions such as the default mode network/ medial frontoparietal network as well as the reward system (e.g., the ventral striatum). Our observation that regions associated with the default mode network are engaged in response to positive emotions during affective empathy is also in accordance with previous studies (Ziaei et al., 2019; for a review see Morelli et al., 2015). Based on our results, we propose that there might be a distinction in the brain networks recruited by older adults in response to positive vs. negative emotions, as a function of task context and task effort required; a hypothesis that needs to be followed up in future studies.

We furthermore provide first evidence here of posterior cingulate bundle involvement during affective empathy for positive emotions in both younger and older adults. In particular, we saw a high concordance between regions that were connected by the posterior cingulum and regions that were activated for positive empathy during affective empathy. A growing body of work now supports that the default mode network might play a role in the processing of positive emotions, possibly due to lower cognitive resources involved in processing of positive (relative to negative) stimuli, and greater salient features of positive cues (e.g., showing teeth; Ziaei & Fischer, 2016). Additionally, research has previously shown that the regions connected with the cingulum bundle have a high overlap with regions within the default mode network (van den Heuvel et al., 2008). However, what had not been demonstrated before, and our results speak to this gap, is the role of these areas for affective empathy, and especially for affective empathy in response to positive emotions in aging. Older adults with higher FA values in the posterior cingulum bundle tract exhibited higher activity in this network for positive emotions. This result again aligns with the motivational account of empathy and provides first evidence of a structure-function link in affective empathic responding to positive emotions in aging.

The activation of the posterior cingulate during positive affective empathy may reflect self-referential (affective) processing in linking one’s own and another’s emotional state to enable an adequate empathic response. In support of this interpretation, the posterior cingulate cortex has been shown to play a role in a wide range of social-cognitive processes (Brewer et al., 2013; Sperduti et al., 2012). For example, posterior cingulate activation is involved in theory of mind and mentalizing (Frith & Frith, 2006; Mitchell, 2009; Molenberghs et al., 2016). A meta-analysis furthermore showed that the posterior cingulate cortex subserves empathy (Bzdok et al., 2012a), specifically the evaluation of how “one relates to one’s experience” (Brewer et al. (2013). Our findings importantly add to previous literature by demonstrating a role for the posterior cingulate in affective empathy in aging. Our findings are in line with the last-in-first-out hypothesis (Madden et al, 2019) that proposes that prefrontal cortex areas, relative to posterior parts of the brain, are the first affected by the aging process. Additionally, our results support the posterior-to-anterior shift in aging (Davis et al., 2008), suggesting that in older compared to younger adults, anterior brain regions (e.g., prefrontal cortex) are more activated than posterior brain regions (e.g., sensory and visual cortex) across a variety of tasks. The higher FA values in the posterior cingulum associated with affective empathy observed in the present study suggest that structural integrity of this region plays a role in affective empathic response, specifically for positive emotions.

We acknowledge that we considered reaction time in this study as the indicator for performance in cognitive and affective empathy. In real life, adequate empathic response may not be evaluated with how fast we respond, but how appropriate our response is given the nature of the stimuli we respond to, as well as how well we can express our empathic concern towards others. Future studies are needed to integrate behavioral responses such as response time with real-life physiological and behavioral measures to study empathy in aging. Additionally, future research is needed to investigate more ecologically valid ways to assess empathic response to emotional stimuli and to investigate the relationship between psychiatric symptoms, such as depression and anxiety, and empathic response among older adults.

## Conclusion

This is the first study to combine behavior with structural and functional brain measures in the study of cognitive and affective empathy towards positive vs. negative emotions among younger and older adults. Older (but not young) adults engaged the salience network during cognitive empathy in response to negative emotions, which could reflect their difficulty in the processing of and/or their enhanced interoception for negative emotions during cognitive empathic responding. In contrast, during affective empathy towards positive emotions, younger and older adults comparably recruited a bilateral network that included nodes of the default mode network; possibly reflecting self-referential processing and/or decreased cognitive effort during affective empathic response to positive emotions. Finally, white matter microstructure of the posterior cingulum bundle was related to positive affective empathy, suggesting that microstructural integrity provides structural support for the functional networks engaged during positive affective empathy.

Findings from this work show that valence plays a critical role in empathic response both in younger and older adults and therefore needs to be considered in investigations into higher-order social-cognitive functions not only in the field of gerontology but also in other populations with deficits in these domains (e.g., individuals with autism spectrum disorder, neurodegenerative disease (Henry et al., 2016), or epilepsy (Ziaei et al., 2021)(Ziaei et al., 2021)(Ziaei et al., 2021). Our results can inform future investigation into the extent to which emotions displayed by another affect social interactions such as closeness or altruistic behavior as well as general well-being via empathic responses, both in young and older adulthood.

## Data and code availability statement

The data is available from this link on OSF repository (https://osf.io/m8zgw/)

## Acknowledgment

The authors would like to thank the participants for their time and acknowledge the practical support provided by the imaging staff at the Centre for Advanced Imaging. We would also like to thank Megan Campbell and Nicola Pease for their help during the data collection. This work was supported by New Staff Research Funding from the Centre for Advanced Imaging and data were collected at the Centre for Advanced Imaging. The authors declare no competing financial interests.

## Footnotes

1 Behavioral performance was analyzed using the three DASS-21 subscales and personal distress subscale from the IRI as covariates. Inclusion of these covariates did not change the results.

2 For completeness, however, results pertaining to accuracy are reported in the Supplemental Material.

3 We have re-run whole-brain analyses with 1000 resampling iterations and results did not change.

